# Potent GST ketosteroid isomerase activity relevant to ecdysteroidogenesis in the malaria vector *Anopheles gambiae*

**DOI:** 10.1101/2023.03.01.530595

**Authors:** Yaman Musdal, Aram Ismail, Birgitta Sjödin, Bengt Mannervik

## Abstract

Nobo is a glutathione transferase (GST) crucially contributing to ecdysteroid biosynthesis in insects of the orders *Diptera* and *Lepidoptera*. Ecdysone is a vital steroid hormone in insects, which governs larval molting and metamorphosis, and suppression of its synthesis has potential as a novel approach to insect growth regulation and combatting vectors of disease. In general, GSTs catalyze detoxication, whereas the specific function of Nobo in ecdysteroidogenesis is unknown. We report that Nobo from the malaria-spreading mosquito *Anopheles gambiae* is a highly efficient ketosteroid isomerase catalyzing double-bond isomerization in the steroids 5-androsten-3,17-dione and 5-pregnen-3,20-dione. These mammalian ketosteroids are unknown in mosquitoes, but the discovered prominent catalytic activity with these compounds suggests that the unknown Nobo substrate in insects has a ketosteroid functionality. Nobo Asp111 is essential for activity with the steroids, but not for conventional GST substrates. Further characterization of Nobo may guide the development of new insecticides to prevent malaria.

## Introduction

The glutathione transferase (GST) enzyme Nobo is essential to the biosynthesis of the molting hormone ecdysone in the mosquito *Anopheles gambiae*. The mosquito is a prominent vector of *Plasmodium falciparum*, the parasite causing malaria, and several hundred thousand people die in malaria every year. The disease is spread by bites of parasite-carrying mosquitoes thereby threatening billions of people in >100 countries world-wide. Transmission of the infectious agent is generally prevented by control of the vector by insecticides, mosquito nets, and elimination of breeding sites (WHO 2021). However, the spread of insecticide resistance has led to general agreement that malaria control is in urgent need of novel agents (Ortelli et al. 2003).

Proliferation of mosquitoes is critically dependent on the steroid hormone ecdysone, and a GST enzyme called Nobo (encoded by the gene *Noppera-bo*) was first discovered in the fruit fly *Drosophila melanogaster* to be essential for ecdysteroid biosynthesis (Chanut Delalande et al. 2014; Enya et al. 2014). Nobo is expressed abundantly in the major ecdysone-producing tissues, primarily in the prothoracic gland, but also in testis and ovary (Enya et al. 2014; Enya et al. 2015). Nobo knock-out *D. melanogaster* present with embryonic lethality and a naked cuticle structure; phenotypes typical for mutants showing embryonic ecdysteroid deficiency. Furthermore, Nobo knock-down larvae displayed lowered titers of the ultimate hormone 20-hydroxyecdysone. A corresponding Nobo GST has been discovered also in the mosquitoes *Aedes aegypti and Culex quinquefasciatus* (Enya et al. 2014), which like *An. gambiae* are important vectors of infectious diseases.

A flavonoid inhibitor of Nobo from *Ae. aegypti* was recently demonstrated to have larvicidal activity, thus supporting the notion that Nobo inhibitors could find use as novel insecticides (Inaba et al. 2022). Notably, Nobo is present only in dipteran and lepidopteran insects and not in other life forms including humans (Enya et al. 2014). From this perspective the enzyme is a perfect new target for new preventative and selective insecticides, avoiding collateral toxicity to honey bees and other valuable biological species. In spite of its demonstrated importance for ecdysteroid biosynthesis, the biochemical function of Nobo is unknown. We have initiated investigations of Nobo from the mosquito *An. gambiae*, the major vector of the pathogen *P. falciparum*. A better understanding of the enzymatic properties can form the basis for discovery and design of potent and selective Nobo inhibitors for use in the combating of the malaria vector.

## Results

A gene encoding the 217 aminoacid sequence of Nobo from *An. gambiae* (also named AgaGSTE8) was synthesized and ligated into a plasmid for expression in *Escherichia coli*. The enzyme was purified by Ni-IMAC and was estimated as >95% homogeneous by SDS-PAGE. Gel filtration verified the purity of Nobo and showed the molecular mass of approximately 50 kDa expected for a dimeric GST protein.

The catalytic activity of the purified enzyme was assayed with standard GST substrates (Mannervik and Jemth 2001): two aryl halides, 1-chloro-2,4-dinitribenzene (CDNB) and 1,2-dichloro-4-nitrobenzene (DCNB); cumene hydroperoxide (CuOOH); and two isothiocyanates, phenethyl isothiocyanate (PEITC) and allyl isothiocyanate (AITC). The specific activities with these cnventional substrates varied between 0.4 and 25 µmol/min per mg protein in the range common to many GST enzymes (Table 1).

**Table 1.**
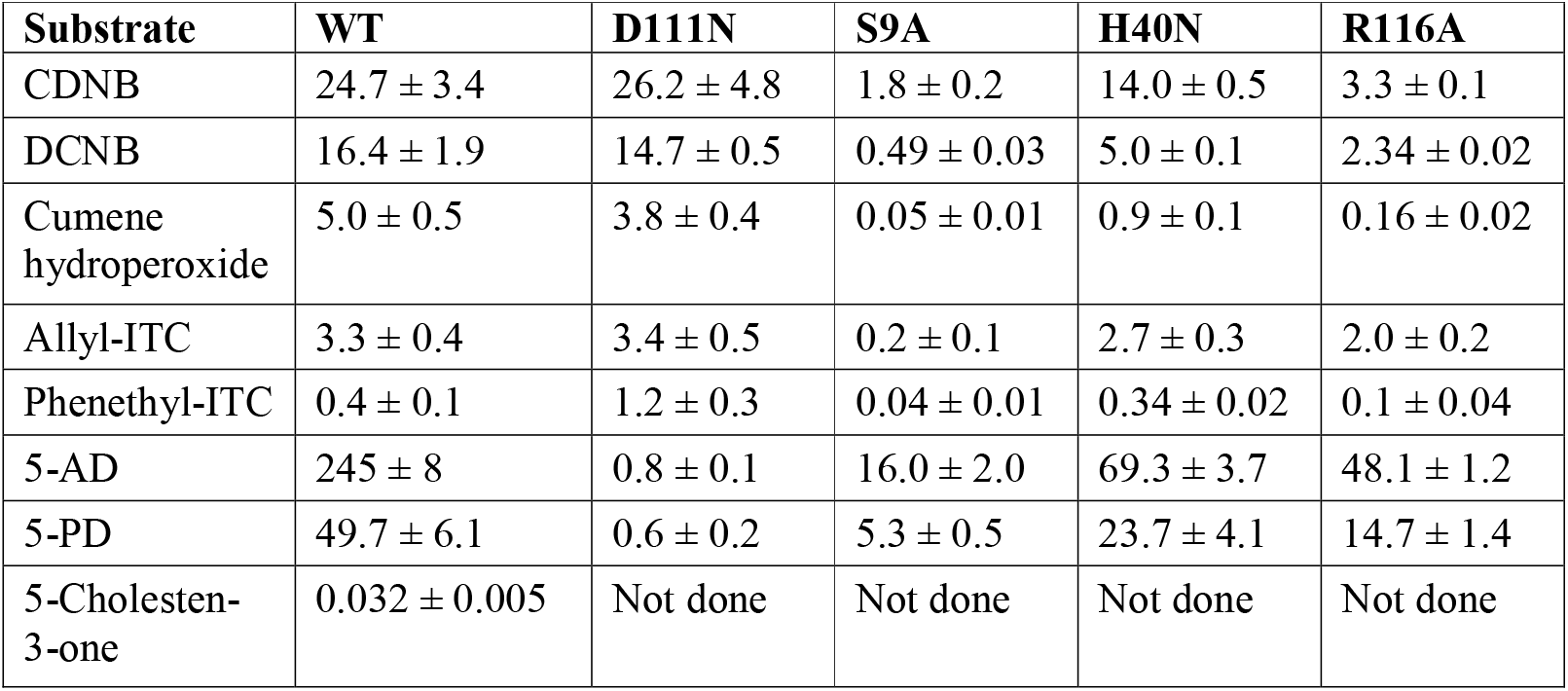
Specific activities (µmol/min per mg) of Nobo from *An. gambiae* with alternative GST substrates. Initial reaction rates at 30 °C were determined spectrophotometrically by published procedures. The values are based on triplicate measurements.

Remarkably, the double-bond isomerase activity with 5-androsten-3,17-dione (5-AD) was 245 µmol/min per mg, significantly higher than the specific activity of any other substrate (Table 1). Similar efficient ketosteroid isomerase catalysis was demonstrated with the alternative steroid substrate 5-pregnen-3,20-dione (5-PD), but the activity with the cholesterol derivative 5-cholesten-3-one was orders of magnitude lower (Table 1). Glutathione was an essential cofactor, although not consumed in the steroid reactions, as earlier demonstrated for mammalian GSTs (Pettersson and Mannervik 2001). Previous studies have identified high double-bond isomerase activity in human (Johansson and Mannervik 2001) and equine GST A3-3 (Lindström et al. 2018), indicating the involvement of GST A3-3 in steroid hormone biosynthesis (Raffalli-Mathieu et al. 2008). A steady-state kinetic analysis of Nobo with 5-AD as the varied substrate determined k_cat_ as 291 ± 6 s^-1^ and k_cat_/K_m_ as (6.9 ± 0.3) · 10^6^ M^-1^s^-1^.

A model of the *An. gambiae* Nobo protein based on the published Nobo structure from *Ae. aegypti* (Inaba et al. 2022) suggested four residues of possible importance for catalytic activity. The following single-point mutations were constructed: D111N, S9A, H40N, and R116A. In D111N the acidic Asp is replaced with the isosteric Asn, and in H40N His is similarly changed to Asn, which cannot act as an acid-base catalyst. The substitution S9A removes the hydroxyl group of Ser, supposedly forming a hydrogen bond to the sulfur of glutathione bound to the active site (Inaba et al. 2022). In R116A removal of the positively charged guanidinium group of Arg is preventing the formation of an ionic bond with a carboxyl group of glutathione. All mutant proteins expressed appeared properly folded and the purified mutant enzymes were tested with the alternative substrates (Table 1).

Notably, the D111N mutation did not markedly change the specific activities with the conventional GST substrates. However, the ketosteroid isomerase activity was dramatically diminished by two orders of magnitude with both 5-AD and 5-PD. The mutation S9A decreased the activity with most substrates by 10 to 20-fold, but 100-fold with CuOOH (Table 1). The mutations H40N and R116A clearly reduced the activities with the ketosteroids, aryl halides, and with CuOOH, but had only a small effect on the isothiocyanate reactions.

In previous studies on Nobo from *D. melanogaster* and *Ae. aegypti* 17β-estradiol was found to be an inhibitor of the enzyme, demonstrating an affinity for steroids (Koiwai et al. 2020; Inaba et al. 2022). The Nobo from *An. gambiae* was likewise inhibited by estradiol with an IC_50_ value 0.17± 0.03 µM. Notably,the D111N mutant was less sensitive to the inhibitor showing an IC_50_ value of 15.5 ± 6.0 µM (Fig. 1).

**Fig. 1.**
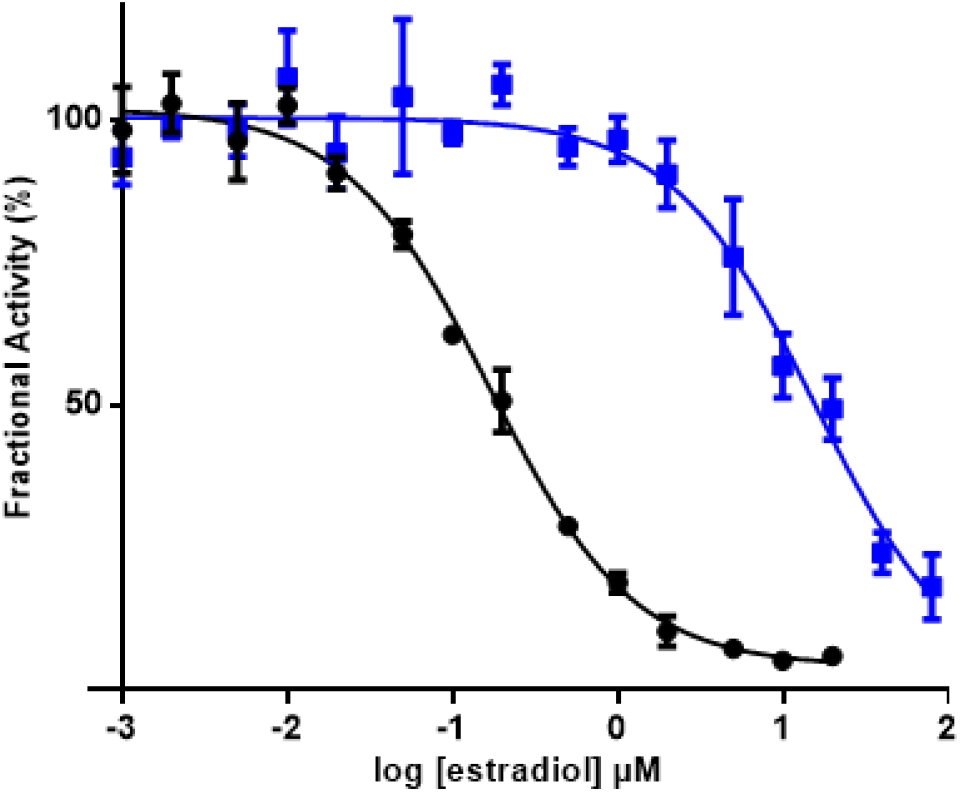
Inhibition of Nobo from An. gambiae with 17β-estradiol. Comparison of wild-type (black dots) with mutant Asp111Asn (blue squares) enzyme using 1 mM DCNB and 5 mM glutathione as substrates and varied 17β-estradiol concentrations at pH 7.5 and 30 °C. Measurements were carried out in triplicate.

### Nobo is an efficient ketosteroid isomerase

The bacterial ketosteroid isomerase is among the most efficient of all enzymes known, with a specific activity of the *Pseudomonas testosterone* enzyme reported as 45,300 µmol/min per mg and the catalytic efficiency k_cat_/K_m_ 1.58 · 10^8^ M^-1^s^-1^ (Kuliopulos et al. 1989), approaching the diffusion limit of enzyme reactions (Pollack 2004). The corresponding values for *An. gambiae* Nobo, 245µmol/min per mg and 6.9 · 10^6^ M^-1^s^-1^, are approximately 20-fold lower, but the mosquito enzyme still qualifies for the echelon of highly active enzymes (Radzicka and Wolfenden, 1995). The requisite in Nobo of the obligatory glutathione in the active site may limit the catalytic turnover as compared with the bacterial ketosteroid isomerase, which is lacking a cofactor. The steroids 5-AD and 5-PD shown to be excellent substrates of Nobo are not intermediates in insect ecdysteroidogenesis, but by mechanistic analogy their efficient isomerization suggests that the enzyme is catalyzing a similar reaction in the “black box” of ecdysone biosynthesis (Pan et al., 2021). The finding that glutathione is required for ecdysteroidogenesis (Enya et al. 2017) lends further support to the notion that the glutathione-dependent Nobo fulfills a catalytic role in the biological context.

Homology modeling of *An. gambiae* Nobo based on the crystal structure of the 51% sequence identical enzyme from *Ae. aegypti* (Inaba et al. 2022) demonstrates that 5-AD (or 5-PD) can fit in an active site pocket apparently reaching the carboxyl group of Asp111. A corresponding carboxylate (Asp113 in *D. melanogaster* and Glu113 in *Ae. aegypti*) is considered a salient and conserved signature feature of all 21 identified Nobo GSTs in *Diptera* and *Lepidoptera* (Koiwai et al. 2020). In the *D. melanogaster* Nobo binding of 17β-estradiol occurs via its 3-hydroxyl group to Asp113 in the conserved site (Koiwai et al. 2020), but binding affinity is lost upon mutation Asp113 into Ala, which eliminates the carboxylate responsible for hydrogen bonding. We propose that Asp111 in *An. gambiae* Nobo makes a bond to the 3-keto group of the ketosteroid substrate, thereby promoting the formation of a dienolate intermediate (Fig. 2). The thiolate group of glutathione bound to Nobo is stabilized by hydrogen bonding to Ser9, like in other GSTs featuring Ser in the active site (Atkinson and Babbitt 2009). The activated thiolate close to the steroid in the active site could serve as a base and abstract a proton from the C4 position, as previously shown in the mammalian GST A3-3 (Dourado et al. 2014).

**Fig. 2.**
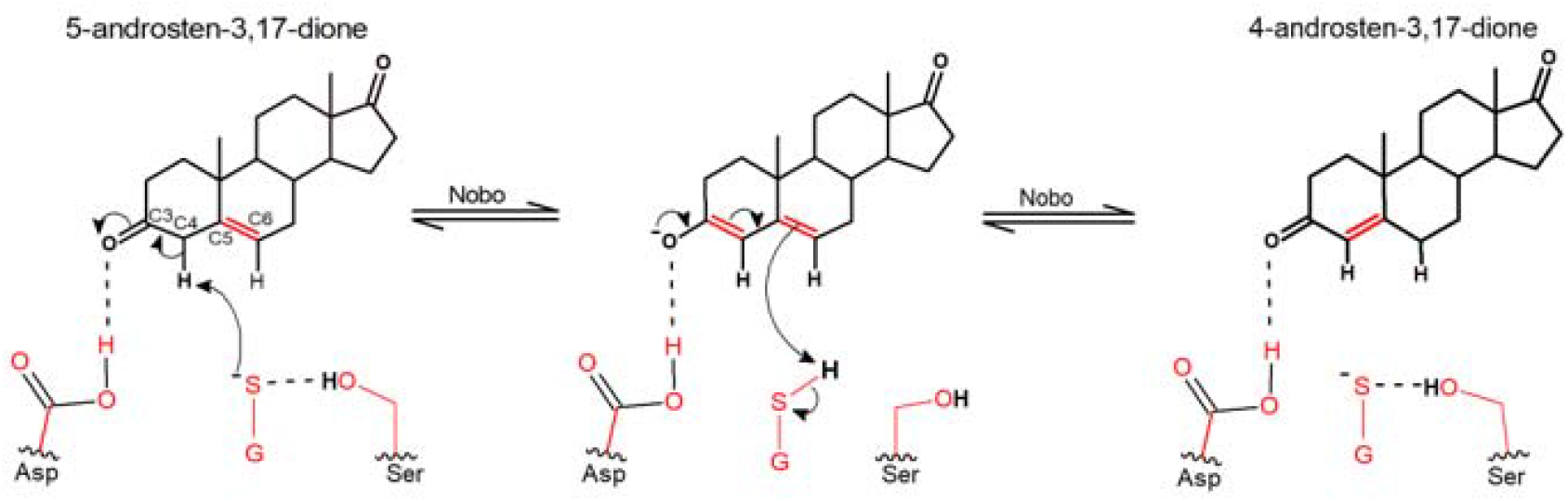
Proposed reaction mechanism of Nobo from *An. gambiae* in the double-bond isomerization of 5-androsten-3,17-dione. Asp111 in a steroid-binding pocket is polarizing the carbonyl group in the C3 position of the substrate. Ser9 is stabilizing the thiolate of glutathione (GSH), which serves as a base abstracting a proton from the C4 position.

The reaction is then completed by addition of a proton to C6 in the steroid. The loss of ketosteroid isomerase activity effected by mutations D111N and S9A in *An. gambiae* Nobo are consistent with these functions (Table 1). The modeled structure indicates that His40 could contribute to binding of glutathione by interacting with the C-terminal carboxylate group of the tripeptide. Arg116 is located at the other side of the same carboxylate and near the orifice of the proposed binding site of the steroid substrate, and like His40 distant from the residues directly involved in catalysis. Thus, the H40N and R116A mutations only modestly diminish the catalytic activity.

Our proposed mechanism for Nobo in the steroid double-bond isomerization is similar to the bacterial ketosteroid isomerase reaction by invoking a carboxylate, which polarizes C=O in position C3 of the steroid substrate (Yabukarski et al. 2022). Like in the mammalian GSTs with steroid isomerase activity (Mannervik et al. 2021), the glutathione thiolate in Nobo serves as the base attacking the C4 hydrogen. In other respects the enzymes differ in functional groups even though they catalyze the same reaction. The primary structures of Nobo proteins and the mammalian GSTs segregate into widely separated evolutionary clades (Koiwai et al. 2020), and neither of them shows any structural relationship to the bacterial ketosteroid isomerase.

## Discussion

Every year more than half a million people die of malaria and other diseases whose pathogenic agents are spread by mosquito bites. The severest form of human malaria is caused by *P. falciparum*, which is transmitted by mosquitoes of the *Anopheles* genus, and the disease is endemic in 76 countries. Transmission of pathogens is largely prevented by control of vector populations by sprayed insecticides, impregnated mosquito nets, and elimination of breeding sites. Malaria control relies heavily on insecticides via indoor residual spraying and long-lasting insecticidal bed nets targeting the principal African vector, *Anopheles gambiae*. However, both resistance and climate change threaten to reverse the progress made by insecticidal mosquito control in recent years. Disappointingly, WHO reports that resistance of the *Anopheles* vectors has emerged to the four main insecticide classes used for mosquito control, i.e. pyrethroids, organochlorines, carbamates, and organophosphates (WHO 2021). Malaria vectors are able to resist the actions of insecticides due to various resistance mechanisms including target site mutations, cuticular modification and metabolic inactivation. The latter occurs by the selection of mosquitoes with upregulated or more efficient endogenous insecticide-detoxifying enzymes. Obviously, novel means to combat the *Anopheles* vectors transmitting the deadly pathogen are in urgent demand (Ranson & Lissenden, 2016).

GSTs are ubiquitous and abundant proteins performing a wide range of enzymatic and non-enzymatic functions (Josephy & Mannervik 2006). In insects, GSTs form an important detoxication system comprised of numerous enzymes involved in the metabolism of a wide range of foreign and endogenous compounds (Koirala et al. 2022). GSTs play an essential role in insect herbivory through the detoxification of deterrent and toxic plant allelochemicals. Research on GSTs in insects was initially motivated by their plausible involvement in insecticide resistance (Ketterman et al. 2011), since elevated GST activity had been detected in strains of insects that are resistant to organophosphates and organochlorines (Adolfi et al. 2019). Insect GSTs have been divided into six classes of cytosolic enzymes: Delta, Epsilon, Omega, Sigma, Theta, and Zeta (Ranson and Hemingway 2005). In addition, membrane-bound insect GSTs exist like the mammalian MAPEG proteins (Bresell et al. 2005). Members of the Delta and Epsilon-class GSTs have been associated with detoxication of various chemicals, including DDT, an insecticide used in indoor residual spraying. The *An. gambiae* GST Nobo in the present investigation (also called GSTE8, Ortelli et al. 2003) belongs to the Epsilon class and its orthologs in *D. melanogaster* and *Ae. aegypti* are designated GSTE14 and GSTE8, respectively. It is noteworthy that neither Delta nor Epsilon GSTs are present in humans and other mammals or in plants.

Ecdysone is a steroid that serves as a master regulator in the control of development, involving molting, metamorphosis, and diapause in ecdysozoan animals, which encompass insects and nematodes (Pan et al. 2021). Following its synthesis, the ecdysone molecule is distributed to various tissues, where it is hydroxylated to form 20-hydroxyecdysone. Transcriptional gene regulation is effected by binding of 20-hydroxyecdysone to a cognate nuclear receptor in target tissues. The biosynthesis of ecdysone originates in dietary cholesterol, and the accepted biosynthetic pathway is initiated by 7,8-dehydrogenation of cholesterol and is terminated by a series of reactions catalyzed by different cytochrome P450 (Halloween) enzymes leading to ecdysone and finally 20-hydroxyecdysone. In between, a number of chemical transformations take place, which remain partly unknown (Black Box in Fig. 3).

**Fig. 3.**
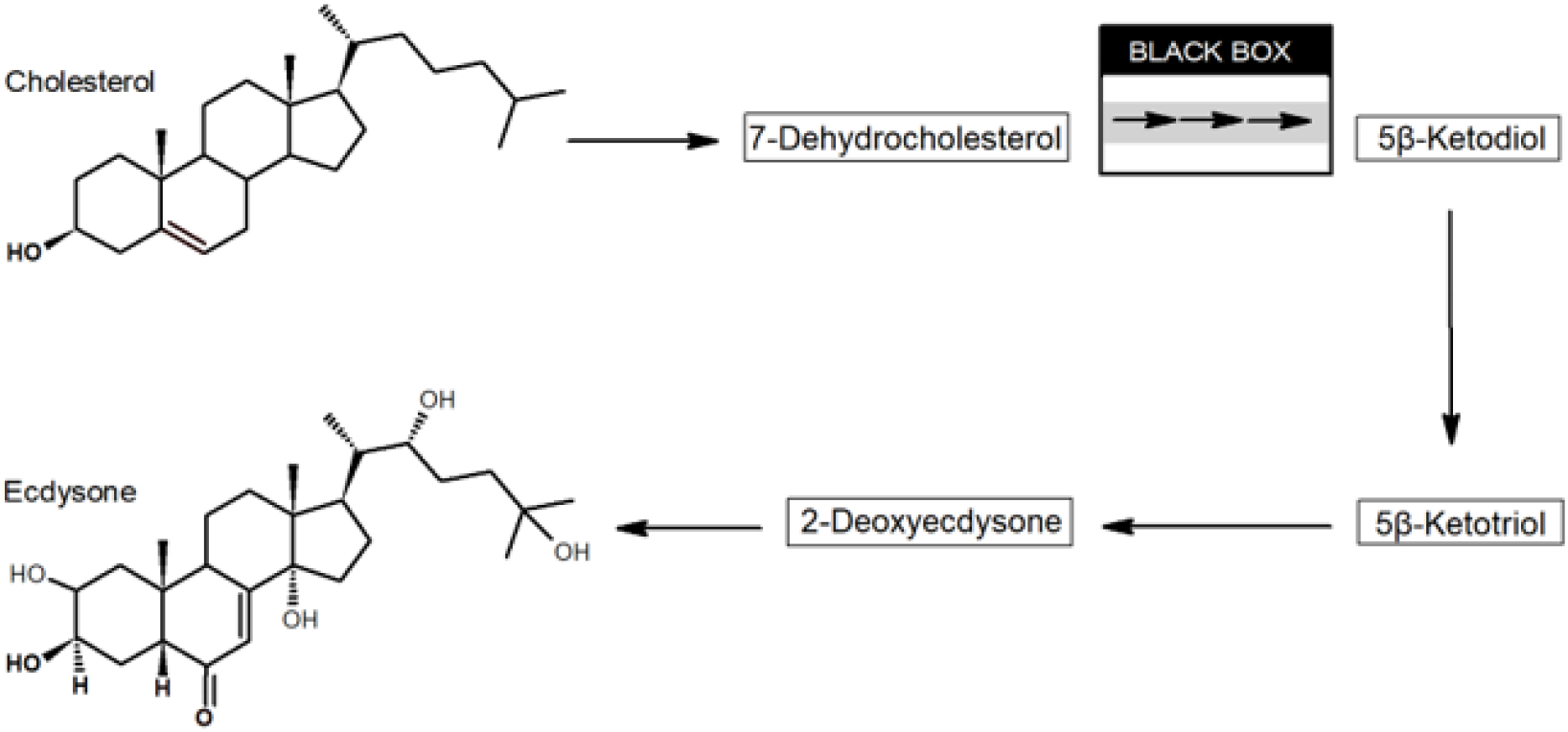
Biosynthetic pathway leading from cholesterol to the hormone ecdysone in insects. Established intermediates are indicated in boxes, whereas the Black Box includes several reactions remaining undisclosed (Warren et al. 2009).

Following dehydrogenation of cholesterol leading to 7-dehydrocholesterol (7dC) it has been hypothesized that reactions in the Black Box involve oxidation of the C3 hydroxyl group in 7dC to yield 3-oxo-7dC (cholesta-5,7-diene-3-one), which may undergo double-bond isomerization to cholesta-4,7-diene-3-one (Warren et al. 2009). Both 3-oxo-7dC isomers are unstable, but it has been demonstrated that photosensitive, ketone-blocked derivatives release 3-oxo-7dC upon irradiation with innocuous UV-light. Administered in vivo by this photolytic approach, 3-oxo-7dC is efficiently incorporated into ecdysteroids by both *D. melanogaster* and the tobacco hornworm moth *Manduca sexta* (Warren et al. 2009). The three reactions leading from 5β-ketodiol to ecdysone are catalyzed by known cytochrome P450 enzymes present in the prothoracic gland. Ecdysone is released into the hemolymph and oxidized into 20-hydroxyecdysterone in peripheral tissues (Kamiyama and Niwa 2022).

In 2014 two research groups identified the gene product of *Noppera-bo* as GSTE14 in *D. melanogaster* (Enya et al. 2014; Chanut-Delalande et al. 2014). Like Halloween cytochrome P450 gene knockouts, null alleles of *Noppera-bo* result in embryonic lethality, embryonic cuticle abnormalities, and decreased ecdysteroid concentrations. Curiously, administration of either cholesterol or 20-hydroxyecdysterone rescues the phenotype, leaving the role of the enzyme Nobo (GSTE14) and the effect of knock-down of the gene unexplained. Niwa and coworkers have identified orthologous GSTs of the Epsilon class in numerous insects, but limited to the orders *Diptera* and *Lepidoptera* (Koiwai et al. 2020).

In addition to the well-established roles of ecdysteroids in metamorphosis and molting, recent investigations further demonstrate the crucial importance of ecdysone signaling for the proliferation of insects (Swevers 2019). For mosquitoes in particular, both oogenesis and larval development are critically dependent on ecdysone in both *An. gambiae* and *Ae. aegypti* (Roy et al. 2018). Genetic modification of *An. gambiae* has demonstrated that disruption of the ecdysone pathway is associated with embryonic lethality midway through development (Poulton et al. 2021). Furthermore, a recent study of mosquito larvae demonstrated that larvicidal activity was correlated with inhibition of Nobo (Inaba et al. 2022). Since ecdysteroids are essential to molting and proliferation in insects, their biosynthesis is therefore a possible target for insect growth regulation and pest control.

The spread of insecticide resistance in *Anopheles* species has prompted the search for alternative sustainable approaches to control malaria. Pyrethroid resistance is, according to the WHO, the biggest biological threat to malaria control. Although new bed nets containing pyrethroid synergists or additional insecticides are slowly being introduced to tackle the loss of insecticidal activity of existing pyrethroid-only nets against increasingly resistance mosquitoes (Grisales et al. 2021), it is clear that malaria control is in dire need of novel agents (Ranson et al. 2016).

The structure of Nobo from *D. melanogaster* has been solved by our group (Škerlová et al. 2020) and by Koiwai et al. (2020), and the enzyme was found to interact with steroid molecules similar to the insect hormone ecdysone and its precursor cholesterol. We discovered that the *D. melanogaster* enzyme catalyzes a ketosteroid double-bond isomerization of 5-androsten-3,17-dione, but the catalytic efficiency was 20-fold lower than that of Nobo from *An. gambiae* reported here.

The GST Nobo enzyme is fundamental to the proliferation of dipteran and lepidopteran insects, being crucial for metamorphosis and oogenesis (Baldini et al. 2013), and inhibitors of the enzyme can be expected to have similar lethal effects as demonstrated in homozygous *nobo* null and Asp113A mutants in *D. melanogaster* (Koiwai et al. 2020). These deleterious effects on ecdysteroidogenesis are noted by Nobo knockout or inhibition in a range of distantly related insect species. For example, similar lethal phenotypes are observed in *Bombyx mori* Nobo null insects (Enya et al. 2015). In mosquitoes, inhibitors of the Nobo ortholog from the yellow-fever transmitting mosquito *Ae. aegypti* are larvicidal to this insect and suppress the expression of the ecdysone-inducible gene *E74B* (Inaba et al. 2022).

Globally the number of malaria cases has stayed in excess of 200 million every year since 2000, but the tally of deaths has declined from 738,000 in the year 2000 to 409,000 in 2019. However, the number of fatalities has reportedly risen to 619,000 in 2021 according to WHO. In spite of some reduced mortality in this period, the rate of improvement has stalled in many countries due to emerging resistance of mosquito vectors to preventative measures. There is general agreement that targeting the insects transmitting the infectious agent remains the mainstay in combating malaria, and novel insect growth regulators are urgently needed to supplement or replace pyrethroids and other insecticides currently used. Incisive studies of Nobo may pave the way to new agents to combat malaria by targeting the enzyme.

## Methods

### Materials

The steroids 5-androsten-3,17-dione and 4-androsten-3,17-dione were purchased from Steraloids Inc. (Newport, RI, USA). All other chemicals were obtained from Sigma-Aldrich (St. Louis, MO, USA).

### Extracting plasmid DNA from filters

DNA encoding Nobo from *An. gambiae* (also known as GSTE8) was synthesized by ATUM using codons promoting high-level expression in *E. coli* (Newark, CA, USA). The gene was ligated into the pD444-SR expression plasmid and delivered on a GF/C glass microfiber filter. Site-directed mutants of Nobo were similarly produced by chemical synthesis. The DNA was extracted with 100 μL 10 mM Tris-HCl, pH 7.5, and centrifuged for 1 min at 10 000×g, yielding 90 μL containing the extracted DNA at a concentration of 20 ng/μL.

### Transformation of bacteria

*E. coli* BL21 (DE3) cells were transformed with 2 μL (40 ng) of the extracted DNA. The cells were kept on ice for 30 min before being heat shocked in a 42 °C water bath for 45 s and then put on ice for 2 min. LB broth (250 μL) was added and the bacteria were grown at 37 °C with shaking for 45 min. The transformed cells were plated on agar with ampicillin (50 μg/mL) and incubated overnight at 37 °C.

### Expression and purification of recombinant GSTs

An overnight culture was prepared from a colony of transformed bacteria suspended in 50 mL LB medium containing ampicillin (50 μg/mL) and incubated overnight at 37 °C with 200 rpm shaking. A flask of 500 mL expression medium (2-TY, 8 g bacto-tryptone, 5 g yeast extract, 2.5 g NaCl, and 25 mg ampicillin) was inoculated with 5 mL overnight culture and grown at 37 °C, 200 rpm. When OD_600_ reached 0.4 enzyme expression was induced with 0.2 mM isopropyl β-D-1-thiogalactopyranoside and the bacteria were further grown for 3 h at 37 °C. The culture was then centrifuged for 7 min at 7000×g. The bacterial pellet was mixed with 10 mL lysis buffer (20 mM sodium phosphate, 20 mM imidazole, 0.5 M NaCl pH 7.4, 0.2 mg/mL lysozyme, and one complete mini tablet of protease inhibitor cocktail) and incubated for 1h. Finally, the cells were disrupted by sonication and the lysate was centrifuged for 30 min at 27,000×g.

The GSTs contained an N-terminal hexahistidine tag enabling their purification by immobilized metal affinity chromatography (IMAC) using a Ni-IMAC column (GE Healthcare). The supernatant fraction from the centrifugation was loaded onto the column equilibrated with binding buffer (20 mM sodium phosphate, 20 mM imidazole, 0.5 M NaCl, pH 7.4), and the column was rinsed with the binding buffer to remove unbound material. An elution buffer (20 mM sodium phosphate, 500 mM NaCl, 250 mM imidazole, pH 7.4) was used to release GST from the column. The eluted GST was dialyzed two times against 10 mM Tris-HCl, pH 7.8, 1 mM EDTA, 0.2 mM tris(2-carboxyethyl)phosphine. SDS-PAGE (sodium dodecyl sulfate-polyacrylamide gel electrophoresis) verified the purity of the dialyzed enzyme.

### Kinetic experiments

Enzyme activities were determined with a selection of standard GST substrates (Mannervik and Danielson 1988). Measurements were performed spectrophotometrically at 30 °C as described in detail previously, including the wavelengths and extinction coefficients used to determine reaction rates (Lindström et al. 2018; Škerlová et al. 2020). However, the assay systems were supplemented with 0.1% (w/v) bovine serum albumin, which gave more reproducible measurements. It should be noted that due to limitations of solubility 5-AD was tested at 0.10 mM whereas 5-PD was limited to 0.010 mM in the determination of specific activities. The inhibition experiments were performed in 96-well plates at pH 7.5 in 0.1 M sodium phosphate buffer with 1.0 mM DCNB, 5.0 mM glutathione, and 17β-estradiol dissolved in ethanol (5% final concentration in the assay system). Kinetic data were evaluated by nonlinear regression analysis using GraphPad Prism 9.

## Acknowledgments

The project was supported by a stipend from Wenner-Gren Foundations for Y.M. and grants to B.M. from the Swedish Research Council. Characterization of Nobo was initiated in our laboratory by Ms. Jessica Khin San and Mr. Kh. Nur E Sadid as students of the University of Skövde.

## Author contributions

B.M. conceived the study and wrote the manuscript with contributions from all other authors. Y.M. performed the kinetic studies; A.I. and B.S. assisted in the enzyme purification and provided materials. All authors contributed to the analysis and discussion of results.

